# Pathogen susceptibility and fitness costs explain variation in immune priming across natural populations of flour beetles

**DOI:** 10.1101/271775

**Authors:** Imroze Khan, Arun Prakash, Deepa Agashe

## Abstract

In many insects, individuals primed with low doses of pathogens live longer after being exposed to the same pathogen later in life. Yet, our understanding of the evolutionary and ecological history of priming of immune response in natural insect populations is limited. Previous work demonstrated population-, sex- and- stage specific variation in the survival benefit of priming response in flour beetles (*Tribolium castaneum*) infected with their natural pathogen *Bacillus thuringiensis*. However, the evolutionary forces responsible for this natural variation remained unclear. Here, we tested whether the strength of the priming response (measured as the survival benefit after priming and subsequent infection relative to unprimed controls) was associated with multiple fitness parameters across 10 flour beetle populations. Our results suggest two major selective pressures that may explain the observed inter-population variation in priming: (A) Basal pathogen susceptibility – populations that were more susceptible to infection produced a stronger priming response, and (B) Reproductive success – populations where primed females produced more offspring had lower survival benefit, suggesting a trade-off between priming response and reproduction. Our work is the first empirical demonstration of multiple selective pressures that may govern the adaptive evolution of immune priming in the wild. We hope that this motivates further experiments to establish the role of pathogen-imposed selection and fitness costs in the evolution of priming in natural insect populations.

## Introduction

Immune priming is now regarded as an integral feature of insect immunity, where exposure to low doses of a pathogen can prime the immune response and confer increased protection against subsequent challenge by the same pathogen (reviewed in Little and Kraaijeveld 2004; Milutinović et al. 2015; Contreras-Garduño et al. 2016). Mathematical models predict that the strength of the priming response can have major implications for infection prevalence, epidemiology and population stability in the wild (Tidbury et al. 2012; Best et al. 2013). Thus, it is important to understand how priming ability evolves in insects and what selective forces drive its evolution in natural populations. In a previous study, we reported considerable variation in the priming response (13-fold survival benefit to no detectable response) among wild-caught populations of the red flour beetle *Tribolium castaneum* (Khan et al. 2016). However, a major unanswered question is – what evolutionary forces lead to such large population level divergence in immune priming?

In general, host immune function can vary due to spatially variable strength of pathogen pressure (Linhart and Grant 1996; N. Reznick and K. Ghalambor 2001; Corby-Harris and Promislow 2008; Mayer et al. 2015). Classic examples of immune investment shaped by parasite-mediated selection include migratory shore birds (Mendes et al. 2006) and island populations of Darwin’s finches (Lindström et al. 2004), in which investment in immune defence (e.g. increased production of natural antibodies) is correlated with infection prevalence. In insects, encapsulation ability increases with the virulence of the pathogen (Kraaijeveld and Alphen 1994), and natural populations of *Drosophila melanogaster* exposed to diverse pathogen communities show an increased ability to clear bacterial infection (Corby-Harris and Promislow 2008). Similarly, strong pathogen selection may also play a direct role in the evolution of the insect priming response (Best et al. 2013), as we demonstrated recently: immune priming evolved rapidly in laboratory selected flour beetle populations exposed to a lethal dose of bacterial infection (Khan et al. 2017). Does pathogen exposure select for priming in natural conditions as well? Wild populations inhabiting pathogen-rich environments have an increased likelihood of reinfection by the same pathogen, and should thus invest more in specific protection against those pathogens (discussed in Corby-Harris and Promislow 2008). We thus reasoned that the strength of the priming response in natural populations can also be determined by the severity of the infection, such that populations with increased pathogen susceptibility should evolve under selection for a stronger priming response.

The maintenance and deployment of immune responses may be costly (Sheldon and Verhulst 1996; Norris and Evans 2000), and it is possible that natural populations vary in their investment in the immune system. If mounting a priming response is metabolically and energetically costly, trade-offs with other fitness components may constrain the strength of immune priming. Mathematical models already highlight the importance of the fitness costs of priming (Tate and Rudolf 2012; Tidbury et al. 2012; Best et al. 2013), although there are only a few studies that experimentally demonstrate costs of priming. For instance, primed female mosquitoes (Contreras-Garduño et al. 2014) and offspring of primed tobacco hornworms (Trauer and Hilker 2013) lay fewer eggs, suggesting a trade-off between priming and reproduction. Maternal immune priming also prolonged offspring development time in mealworm beetles (Zanchi et al. 2011), compromising their competitive ability at high density (Koella and Boete 2016). Based on these results, we speculate that variable fitness costs could also determine the occurrence and maintenance of priming ability in natural populations.

To elucidate the selective parameters underlying variation in priming, we analysed the response of 10 natural populations of the red flour beetle *Tribolium castaneum* primed with the natural pathogen *Bacillus thuringiensis* (previously described in Khan et al 2016). For each of these populations, we measured the within-generation priming response, basal pathogen susceptibility (without priming), and various fitness and immune components. First, we asked whether the benefit of priming increases with susceptibility to infection. Second, we tested whether potential fitness costs of immune priming trade off with its survival benefit. Finally, we also tested whether tradeoffs with other immune responses may explain the observed variation in priming responses. Our experiments were thus explicitly designed to detect relationships between various life-history and ecological parameters that may drive populations divergence in priming ability.

## METHODS

### Generating experimental beetles

We collected 10 natural populations of *Tribolium castaneum* from grain warehouses at different geographical locations throughout India (described in Khan et al. 2016). We reasoned that multiple factors such as individual age, mating, migration history, nutrient, and local environmental factors are likely to influence variability in immune responses across populations. Since it was impossible to account for all these factors, we did not measure immune priming responses on individuals that were directly collected from the grain warehouses. Instead, we maintained them in the laboratory on whole-wheat flour at 34 ◻C for one year before commencing the experiments (Khan et al. 2016). To generate experimental individuals for each population, we allowed 800-1000 individuals to oviposit in 350g of wheat flour for 48h. We then removed the adults and collected female offspring at the pupal stage (after ~3 weeks). We discarded males since handling both sexes simultaneously was logistically challenging. We housed 3 female pupae in 2 ml micro-centrifuge tubes containing 1g flour for 12 days. Separately, we also collected larval offspring after 10 days. Since the pupal stage lasts for 3-4 days and eggs usually hatch in 2 days, we obtained 8-day-old adults and 8-day-old larvae were for all our experiments.

### Immune priming and challenge

We used the natural insect pathogen *Bacillus thuringiensis* to measure within-generation priming for all populations as described in Khan et al (2016). Briefly, we pricked adults between the head and thorax, and larvae between the last and last but one segment, using a 0.1mm insect pin (FST, CA) dipped in heat-killed bacterial slurry (i.e. priming) or insect Ringer (i.e. sham priming control). We made bacterial slurry from 10 ml freshly grown overnight culture of *B. thuringienses* at 30°C (optical density of 0.95; adjusted to ~10^11^ cells in 1 ml insect Ringer solution). We used heat-killed cells to prime beetles because this would induce an immune response without a direct cost of virulence. After this, we isolated experimental beetles (i.e. adults or larvae) in wells of 96-well microplates (Corning) containing wheat flour. Six days later, we checked their mortality and then again challenged individuals with live bacterial culture adjusted to ~10^10^ ml^−1^ (delivering ~900 bacterial cells per beetle). We did not find any mortality after priming and before live pathogen challenge. Control beetles received mock priming followed by a mock challenge with insect Ringer. After the immune challenge, we immediately returned experimental beetles from each population to wells of fresh 96-well micro plates and measured various traits as described below (also see Figure 1). Since we used a low dose of infection (compare with Khan et al. 2016), we observed a late onset of post-infection mortality. For instance, while a few infected larvae (<1%) died before pupation in some populations, there were no deaths during the adult stage until 23 days post-eclosion. We also did not observe any mortality in adults until a week after infection.

**Figure 1.**
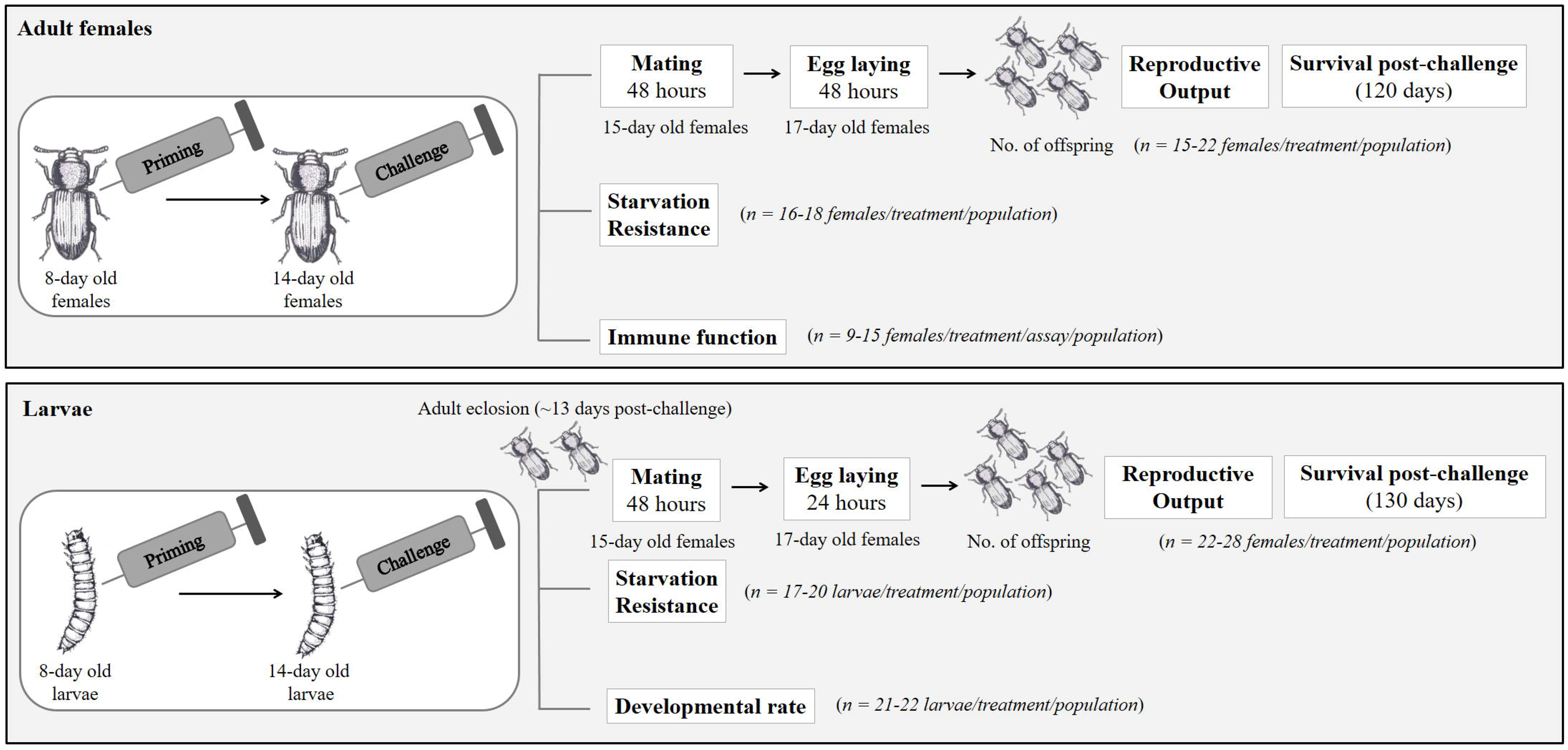
Experimental design.

### Joint assay of basal pathogen susceptibility, survival benefit of priming and changes in reproductive output after priming

One day after the immune challenge, we paired a subset of adult females with uninfected, 8-day-old virgin males from the respective population for 48 hours in a 1.5 ml microcentrifuge tube containing 1 g flour (one pair of beetles per tube). We then separated females to measure the total number of offspring produced by each female (48 h of oviposition; eggs allowed to develop in 6 g flour for 3 weeks). Following this, we returned mated females to 96-well microplates with flour, and noted their survival every 3-5 days for another ~117 days (total 120 days post-challenge), transferring them to fresh microplates with food every 5 days to minimize the interaction between females and new offspring (n=15-22 females/treatment/population). For larvae, we isolated primed and challenged individuals in their respective wells until they pupated. Subsequently, we identified and retained only female pupae. Fifteen days post-eclosion, we paired each adult female with a virgin male as described above. We allowed females to oviposit for 24 hours and recorded their mortality every 3-5 days for another 100 days (total ~130 days post-challenge) as described above (n=22-28 females/treatment/population). This procedure allowed us to obtain a correlated dataset for early reproductive success and survival of each experimental female after priming and challenge. A few replicate plates for reproductive output after adult priming were accidentally lost during the experiment. Hence, the sample size for the individual correlation was lower than expected (See Table S1).

The residuals for reproductive success were not normally distributed (tested with Shapiro-Wilks test). We thus used nonparametric Wilcoxon Rank Sum tests to test the impact of larval and adult priming on reproductive success. We quantified the impact of priming on reproductive output as: Mean number of offspring produced by primed females / Mean number of offspring produced by unprimed females. We analyzed post-immune challenge survival data for each population and life stage separately using Cox proportional hazard survival analysis with priming as a fixed factor and lifespan in days as the response variable. We considered beetles that were still alive at the end of the experiment as censored values. For each population and life stage, we calculated pathogen susceptibility as the estimated hazard ratio of unprimed vs. control groups (Rate of deaths occurring in unprimed group / Rate of deaths occurring in full control group). A hazard ratio significantly greater than one indicates an enhanced risk of mortality in unprimed groups compared to control individuals. To estimate the strength of the priming response, we calculated the survival benefit to the host after infection, with vs. without previous exposure to the same pathogen (Rate of deaths occurring in unprimed group / Rate of deaths occurring in primed group). A hazard ratio significantly greater than one indicates an enhanced risk of mortality in unprimed groups compared to primed individuals.

### Separate assays to measure the impact of priming on development rate, lifespan under starvation and immune components

In these assays, we did not re-measure the immune priming response in terms of survival benefit after infection, since this was already measured for each population as described above. Instead, we directly measured the impact of priming (i.e. compared primed vs. unprimed groups) for the following fitness and immune components.

1. Lifespan under starvation: We first tested whether priming affects lifespan under starvation in different populations. To do this, we isolated a subset of virgin females and larvae individually in 96-well microplates without food (*n =*16-20/lifestage/treatment/population) immediately after immune challenge. We noted their mortality every 12 hours (10 am & 10pm ±1 hour) until all of them died. We quantified the impact of priming (primed vs. unprimed) on lifespan under starvation using hazard ratios, as described above. A hazard ratio significantly greater than one would suggest a higher risk of mortality in primed groups compared to unprimed individuals. Due to logistical reasons, we could only measure starvation resistance of larvae from 7 populations (except B1, AM and ND; described in Khan et al 2016).
2. Developmental rate: To measure the effect of priming on larval development, we placed immune-challenged experimental larvae individually in 96-well microplates (*n =*21-22 larvae/treatment/population). We observed larvae every 12 hours (11 am & 11pm ±1 hour) and noted the time to pupation for each larva. We analyzed the data (non-normally distributed) using nonparametric Wilcoxon Rank Sum tests and calculated the impact of priming on larval development as: Mean time to pupation of primed larvae / Mean time to pupation of unprimed larvae.
3. Immune components: To measure aspects of immune function, we first primed and challenged adult females from each population (described above). Next day, we used a subset of females to quantify antibacterial activity of beetle hemolymph (*n* = 9-15/treatment/population) (see Khan et al. 2015 for detailed methods). Briefly, we measured the zone of inhibition produced by beetle homogenates on a lawn of *B. thuringiensis* growing on nutrient agar medium. Flour beetles also secrete defensive quinone compounds that inhibit the microbial growth in their surroundings, acting as an external immune defense (Joop et al. 2014, Khan et al. 2015). To quantify this immune response, we measured the zone of inhibition produced by cold-shocked females (−86°C for 20 minutes) embedded vertically in a lawn produced by *B. thuringiensis* growth on nutrient agar plates (n = 9-10 females/treatment/population). A cold shock triggers the release of abdominal and thoracic stink gland contents with antimicrobial quinones (Khan et al. 2015). We analyzed the non-normally distributed immune response data using nonparametric Wilcoxon Rank Sum tests and estimated the impact of priming on immune components as: Mean zone of inhibition produced by primed females / Mean zone of inhibition produced by unprimed females.

## RESULTS

The strength of immune priming is usually quantified as the survival benefit to the host after infection, with vs. without previous exposure to the same pathogen (Roth et al. 2010;Khan et al. 2016). Hence, we quantified the strength of the priming response as the proportional hazard ratio estimated from survival data for primed vs. unprimed individuals, all subsequently infected with live bacteria. In most populations, both larval and adult survival increased significantly after priming (adults: 9/10 populations, larvae: 8/10 populations; **Fig. 2A**; see **Figs. S1 & S2** for survival curves). As reported earlier (Khan et al. 2016), the priming response also varied substantially across populations, ranging from no detectable response to a 10-fold increase in larval post-infection survival relative to the unprimed control. We found that this variation was strongly associated with susceptibility to infection, measured as the hazard ratio for infected vs. uninfected groups (**Fig. 2A**). These results are consistent with the prediction that the severity of infection may determine the strength of immune priming in insect populations (Best et al. 2013) – more susceptible populations may face stronger selection for evolving priming response.

**Figure 2.**
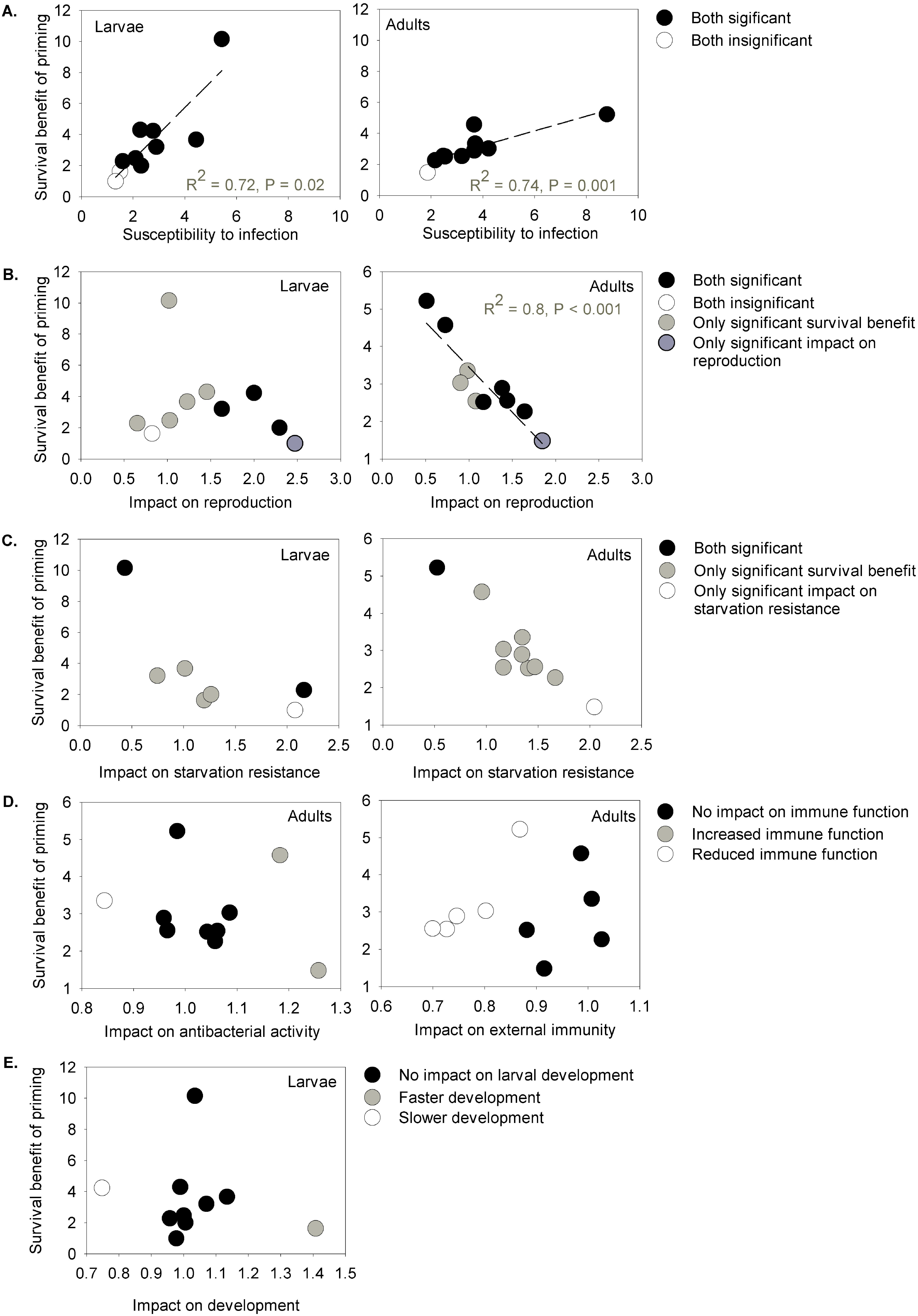
Correlation between the survival benefit of immune priming and (A) basal susceptibility to infection (B) reproductive benefit of immune priming; impact on (C) starvation resistance (D) antibacterial activity (E) external immunity and (F) larval developmental rate.

The costs of mounting a priming response can also vary across natural populations, in turn limiting the expression of priming. To test this hypothesis, we analyzed the impact of priming on multiple fitness-related traits: reproduction, larval development, lifespan under starvation, and other immune components. As predicted, we found a negative correlation between the strength of adult immune priming and subsequent change in reproductive fitness, though there was no such association for larvae (**Fig. 2B**). Contrary to the cost hypothesis, we found that only two populations showed a significant decrease in reproductive fitness after adult priming (i.e. significant reproductive impact <1). In most populations, females either produced more offspring after priming (5/10) or showed no detectable change in reproduction (3/10) (Fig. 2B **& S3**). Similarly, for larval priming, none of the populations showed a significant cost of reproduction (**Fig. S3**). Together, these results suggest that although priming generally does not impose a reproductive cost, it may induce a stage-specific trade-off between survival vs. reproductive benefits.

Overall, priming did not affect starvation lifespan, except for a few populations where it either prolonged (Larvae: 2/7 populations; Adults: 1/10 populations) or reduced lifespan (Larvae: 1/7 populations; Adults: 1/10 populations) (see **Fig. S4 & S5** for survival curves). Although there was no consistent association between larval priming and lifespan under starvation, our data suggest a potential relationship between adult investment in priming vs. starvation lifespan. Individuals from the population that lacked a priming response lived twice as long under starvation, whereas adult beetles from the population showing the strongest priming response (~5-fold survival benefit) died faster under starvation (**Fig. 2C**). These results suggest a possible cost of priming response in terms of depleting energy reserves, in turn reducing lifespan under starvation.

We found that immune priming had a contrasting effect on two components of adult immunity. While priming consistently reduced external immunity in several populations (5/10), its impact on antibacterial activity was highly variable (**Fig. S6**). Priming had no impact on antibacterial activity in most populations (7/10), except a few of them where primed females produced either larger (2/10) or smaller zones of inhibition (1/10) than unprimed controls. None of the immune components was correlated with the observed variation in priming response across populations (**Fig. 2D**).

Finally, we tested whether larval immune priming traded off with larval development rate across populations. We found that immune priming did not alter larval development except in two populations, where larvae either developed faster (AG) or showed delayed development (ND) compared to controls (**Fig. S7**). Thus, developmental rate of primed larvae could not explain population level variation in larval priming response (**Fig. 2F**).

## DISCUSSION

We present the first systematic test of multiple factors that may determine the evolution of the strength of immune priming in natural insect populations. Previously, we documented large variation in the priming response among wild-caught flour beetle populations (Khan et al. 2016), ranging from no detectable response to a 13-fold survival benefit in some populations. Although this work provided an empirical framework to understand whether and to what degree priming responses vary in natural populations, the selective forces responsible for this variability remained unclear. Previous theoretical work suggests that the strength of the priming response may depend on the strength of selection imposed by pathogens (Tate and Rudolf 2012; Best et al. 2013), as well as constraints imposed by the cost of immune priming via tradeoffs with other fitness components. Our results are consistent with both types of selection. First, we show that the bacterial pathogen *B. thuringiensis* has a variable impact in different populations, suggesting that the same pathogen can impose divergent selection pressure across populations of the same host species. Subsequently, we find that increased susceptibility to *B. thuringiensis* is positively correlated with an increased survival benefit of immune priming, such that priming is most beneficial for populations that are highly susceptible to infection. This indicates that the pathogen-mediated reduction in lifespan may act to increase selection for priming response in natural populations. Second, we found a life stage-specific negative relationship (a possible trade-off) with reproductive success that may constrain the strength of priming. When primed and infected adult females produced more offspring than unprimed controls (i.e. priming increased reproduction), they showed a reduced survival benefit of priming. Thus, the most important implication of our work is that both specific fitness costs and the fitness impact of infection can determine variability in priming, as well as reflect conditions that may favor the evolution of stronger priming.

A notable strength of our study is the use of multiple natural populations to gain deeper insights into the evolutionary and ecological history of priming in an insect. Our results indicate a general adaptive role of immune priming such that in many populations, priming not only improves long-term lifespan but also leads to an immediate gain in reproductive effort. This seems to contradict a mathematical model that predicts large reproductive costs associated with priming (Best et al. 2013). However, a careful comparison across populations revealed that immune priming does not improve survival and reproduction equally in a population. Instead, greater benefits of reproduction come at the cost of reduced survival, suggesting a broadly distributed hidden trade-off between these traits. A recent report in mosquitoes also showed that primed females that invested more in egg production show reduced pathogen clearance and greater susceptibility to infection (Contreras-Garduño et al. 2014). Such parallel results from intra-vs. inter-population studies suggest the existence of trade-offs at multiple levels. Our data also suggest a weak negative association between priming and starvation resistance. Although priming had no impact on starvation resistance in most populations, populations where priming maximized survival benefit after infection (i.e. ND) or showed no priming response (i.e. AG) also had respectively reduced or increased starvation resistance. Previous studies have documented trade-offs between immune investment and lifespan during starvation in fruit flies (Valtonen et al. 2010) and bumble bee workers (Moret and Schmid-Hempel 2000). In fruit flies, trade-off between immunity and starvation resistance may also have a genetic basis: genotypes that invest more in immunity have lower survival under starvation and vice versa (Hoang 2001). However, it is currently unclear whether such phenotypic or genetic trade-offs are also widespread with respect to priming ability and lifespan during starvation. Thus, we suggest that our experiments uncover an exciting possibility that requires further work.

What are the mechanisms underlying the potential trade-offs we observed? Both endocrine signaling (e.g. via juvenile hormone and ecdysone) and altered lipid metabolism via insulin/insulin-like growth factor-like signalling pathway (IIS) can mediate physiological trade-offs between immunity and major fitness components (Schwenke et al. 2016; Schwenke and Lazzaro 2017). A previous study in fruit flies suggests that immune activation via the Toll signaling pathway in the fat body (the major immune and lipid storage organ in insects) inhibits insulin signaling activity (DiAngelo et al. 2009). Reduced insulin signalling after infection could further lead to decreased lipid storage (DiAngelo et al. 2009), increased haemolymph lipid concentration (Cheon et al. 2006) and finally, reallocation of energy utilisation to immune response from reproduction (Clancy 2001) or starvation resistance (Beenakkers et al. 1986). Interestingly, recent studies suggest that activation of *Toll* pathways and lipid mobilisation are equally important for mounting a successful immune priming response in fruit flies (Pham et al. 2007) and mosquitoes (Ramirez et al. 2015). This sets up the possibility that similar signalling mechanisms are also responsible for the observed negative relationship between immune priming and fitness components in flour beetles. Further studies may provide greater mechanistic insight into how early reproductive success or starvation resistance can alter later immune priming response.

Another interesting finding of our study is that although priming delayed or accelerated larval development in some populations, it did not explain the observed population level variation in priming response. Previous studies using a single population of different insect species found that immune activation in larvae can accelerate development (Roth and Kurtz 2008) and maternal priming can either accelerate (Tate and Graham 2015) or reduce (Zanchi et al. 2011) offspring development rate. However, ours is the first study that documents the differential impact of immune priming on development rate across populations of the same species. Therefore, an additional implication of our work is that studies using a single population are insufficient to generalize the costs or benefits of priming because priming has variable consequences across populations.

Multiple lines of evidence suggest that priming is beneficial because it induces more efficient immune responses (reviewed in Milutinović et al. 2016). In contrast, our results present a more complicated scenario – priming increased antibacterial activity only in a few populations, and most populations with large survival benefit of priming did not always increase antibacterial activity. Overall, antibacterial activity did not explain the observed variation in primimg response, highlighting that the association between innate immune responses and survival after priming may not be straightforward in wild populations. This is surprising since previous studies with *Tribolium* beetles found that priming with *B. thuringiensis* increases expression of a large set of immune related genes (Greenwood et al. 2017), correlating strongly with survival after reinfection (Milutinović et al. 2013). We note that whereas our hemolymph antibacterial activity assay may reflect the impact of several immune pathways such as antimicrobial peptides, it is possible that other aspects of innate immunity such as cellular defence (e.g. circulating hemocytes) play a more important role in priming in natural populations (Rodrigues et al. 2010).

Finally, we note a novel result where priming exerts opposite effects on different aspects of beetle immunity, e.g. antibacterial activity (internal immunity) of the hemolymph vs. quinone secretion outside the body (external immunity, see Joop et al. 2014). In contrast to its variable impacts on antibacterial activity, priming consistently reduced external immune function in several populations, suggesting a crosstalk between priming and quinone production pathways. Although these results are broadly similar to our previous experiment where bacterial infection reduced quinone production in virgin females (Khan et al. 2015), it did not explain the variation in primimng response across populations. We suggest further manipulative experiments to disentangle the complex interaction between priming, innate immune pathways and quinone production in flour beetles.

Overall, our data highlight the importance of explicitly testing the impact of pathogen selection and fitness costs on the immune system of wild populations. While our results provide valuable insight into the macro-evolutionary patterns of priming evolution, a significant limitation is the lack of information on local pathogen pressure that our beetle populations experienced before we brought them into the laboratory. Recent evidence suggests that *Drosophil*a populations previously exposed to multiple pathogens are more resistant to a novel *Lactobacillus lactis* infection (Corby-Harris and Promislow 2008). Increased resistance to subsequent *L. lactis* infection can also be selectively favored by previous exposure to conspecific *Lactococcus* species. Similarly, it is possible that our beetle populations have already encountered variable selection imposed by widely distributed natural pathogens such as *B. thuringiensis* in their natural habitat. Therefore, the observed responses against experimental manipulations (e.g. priming and/or live pathogen exposure) can also be influenced by previously experienced pathogen selection. Indeed, at the molecular level, many immune-related genes in insects show signs of strong selection, suggesting rapid coevolution with pathogens (Lazzaro et al. 2006; Sackton et al. 2007).

In conclusion, we suggest that our work serves as an important first step towards understanding whether and why natural populations of insects differ in their immune priming respose. Previously, we used experimental evolution to show that strong pathogen selection is necessary to evolve immune priming in laboratory populations of flour beetles (Khan et al. 2017). However, it remains unclear whether wild populations also show a similar response. We suggest future experiments where susceptible wild populations may be allowed to evolve under different levels (low to high) of selection imposed by the pathogen. Such experiments will allow us to directly test for a positive correlation between evolved priming response and the level of pathogen selection, as well as associated evolutionary fitness costs.

## AUTHOR CONTRIBUTIONS

IK conceived experiments; IK, DA and AP designed experiments; AP and IK carried out experiments; IK analyzed data; IK and DA wrote the manuscript with input from AP. All authors gave final approval for publication.

## ACKNOWLEDGEMENTS

We are grateful to Ann Tate for feedback on the manuscript. We thank Swastika Issar and Divya Meena for their help during experiments.

